# Marked changes in diversity and relative activity of picoeukaryotes with depth in the global ocean

**DOI:** 10.1101/552604

**Authors:** Caterina R. Giner, Vanessa Balagué, Massimo C. Pernice, Carlos M. Duarte, Josep M. Gasol, Ramiro Logares, Ramon Massana

## Abstract

Microbial eukaryotes are key components of the ocean plankton. Yet, our understanding of their community composition and activity in different water layers of the ocean is limited, particularly for picoeukaryotes (0.2-3*µ*m cell size). Here we examined the picoeukaryotic communities inhabiting different vertical zones of the tropical and subtropical global ocean: surface, deep chlorophyll maximum, mesopelagic (including the deep scattering layer and minimum oxygen zone) and bathypelagic. Communities were analysed by high-throughput sequencing of the 18S rRNA gene, as represented by DNA (community structure) and RNA (metabolic expression), followed by delineation of Operational Taxonomic Units (OTUs). We found a clear stratification of the picoeukaryotic communities along the water column, with two differentiated assemblages corresponding to the sunlit and dark ocean. Specific taxonomic groups either increased or decreased their abundances with depth. We used the rRNA:rDNA ratio of each individual OTU as a proxy of its metabolic activity. The highest relative activity was found in the mesopelagic layer for most taxonomic groups, and the lowest in the bathypelagic. Overall, our results characterize the change in community structure and activity of picoeukaryotes in the global-ocean water column, suggesting that the mesopeagic layer is a hot-spot of picoeukaryotic activity.

## INTRODUCTION

Protists are key components of marine microbial communities, playing a central role in marine food webs (Sherr and Sherr 2002), particularly in carbon cycling (Arístegui et al 2009). Despite their importance, the distribution of protists in the global ocean is still poorly understood. In particular, little is known about protists communities in the dark ocean, as most efforts have focused in protists populating the photic layer. Yet, the deep ocean (>1,000 m depth) is a huge biome, being the largest reservoir of organic carbon in the ocean (Nagata et al 2010) and containing about 70% of the ocean’s prokaryotic cells (Arístegui et al 2009).

The water column is divided into epipelagic (0-200 m depth), mesopelagic (200-1,000 m) and bathypelagic (1,000-4,000 m) layers. The sunlit epipelagic zone harbors photosynthetic microbes and is the scenario of the classic oceanic food web, whereas the dark ocean (i.e. mesopelagic and bathypelagic regions), are characterized by high pressure, low temperature and high inorganic nutrient content (Arístegui et al 2009). A global survey indicated that the distribution of protists in the upper ocean is predominantly structured by oceanographic basin, pointing to dispersal limitation (de Vargas et al 2015). In contrast, the available evidence on the distribution of protists along the water column is limited to a few specific ocean regions and indicates a clear differentiation between epipelagic and deep ocean communities (Brown et al 2009, Countway et al 2007, Hu et al 2016, Jing et al 2018, López-García et al 2001, Not et al 2007, Xu et al 2017, Xu et al 2018). The mesopelagic layer represents ~20% of the global ocean and is an area of intense remineralization (Arístegui et al 2009), believed to sustain a huge fish biomass (Irigoien et al 2014). It includes a layer few hundred meters thick known as the ‘deep scattering layer’ (DSL), characterized by the accumulation of large stocks of mesopelagic fish and migrant zooplankton (Irigoien et al 2014), leading to intense biological activity. Furthermore, in regions like the eastern tropical Pacific Ocean and the tropical Indian Ocean, the mesopelagic zone contains layers with very low oxygen concentration, known as oxygen minimum zones (OMZ), which play a key role in the nitrogen cycle and contain specific microbial communities (Robinson et al 2010). Protist communities in OMZ or anoxic basins have received less attention than those in oxygenated mesopelagic waters (Jing et al 2015, Orsi et al 2012, Parris et al 2014, Stoeck and Epstein 2003). Furthermore, changes in protist relative activity in the DSL or OMZ at the global scale remain unknown. On the other hand, the bathypelagic layer is assumed to be a highly stable environment, and protist assemblages in this zone appear to be structured by water masses (Pernice et al 2016).

DNA-based analyses have contributed greatly to elucidate global protist community structure in the sunlit (de Vargas et al 2015) and dark ocean (Pernice et al 2016). Yet, sequences obtained using DNA include metabolically inactive or dead cells, while sequences obtained from RNA extracts derive from the actual ribosomes and therefore from living cells. In addition, as the ribosomal content is regulated to match the protein synthesis needs during population growth and acclimation (Blazewicz et al 2013), the rRNA:rDNA ratio for a given taxon has been used as a proxy for their relative activity. This approach has been applied to microbial eukaryotes in a few epipelagic (Hu et al 2016, Massana et al 2015) and vertical-profile (Jing et al 2018, Xu et al 2017) regional studies, but it has not yet been applied at the global scale.

Here we present the first global survey investigating vertical changes in picoeukaryote community composition and relative activity using Illumina sequencing of 18S rRNA genes amplified from DNA and RNA extracts. We analyzed samples from 7 depths (from surface to 4,000 m) in 13 stations encompassing the Atlantic, Indian and Pacific Oceans sampled during the Malaspina-2010 Circumnavigation expedition (Duarte 2015), targeting protists <3 µm in size (picoeukaryotes). Specifically, we analysed changes in picoeukaryote community structure and relative activity along the ocean water column, assessing environmental factors driving these changes. Furthermore, we tested whether picoeukaryote community structure and relative activity differ in the OMZ and DSL zones.

## MATERIALS AND METHODS

### Sample collection and nucleic acid extraction

During the Malaspina-2010 Circumnavigation expedition (December 2010 – July 2011), a total of 91 water samples were collected in 13 stations distributed across the world’s oceans (Fig. S1). Each station was sampled at 7 different depths with Niskin bottles attached to a CTD profiler that had sensors for conductivity, temperature, salinity and oxygen. Each vertical profile included samples at surface (3 m), Deep Chlorophyll Maximum (DCM), and 2-3 depths in mesopelagic (200-1,000 m) and bathypelagic (1,000-4,000 m) waters. Samples for inorganic nutrients (NO_3_-, NO_2_-, PO_4_^3-^, SiO_2_) were collected from the Niskin bottles, kept frozen, and measured spectrophotometrically using an Alliance Evolution II autoanalyzer (Grasshoff et al 1983). Along the cruise, different deep-water masses were sampled. The proportion of the different deep-water masses in each sample was inferred from its temperature, salinity and oxygen concentration (Catalá et al 2015).

For each sample, about 12 liters of seawater were prefiltered through a 200 µm nylon mesh to remove large plankton and then sequentially filtered using a peristaltic pump through a 20 µm nylon mesh and then through 3 µm and 0.2 µm pore-size polycarbonate filters of 142 mm diameter (Isopore, Millipore). Filtration time was about 15-20 minutes. The filters were flash frozen in liquid nitrogen and stored at −80°C until DNA and RNA extraction. Polycarbonate filters containing the 0.2-3 *µ*m size fraction were cut into small pieces and cryo-grinded with a Freezer-Mill 6770 (Spex) using 3 cycles of 1 minute. Then, RNA and DNA were extracted simultaneously using the Nucleospin RNA kit (Macherey-Nagel) plus the NucleoSpin RNA/DNA Buffer Set (Macherey-Nagel) procedures. The presence of residual DNA in RNA extracts was checked by PCR with universal eukaryotic primers and, if detected, was removed using the Turbo DNA-free kit (Applied Biosystems). RNA was reverse transcribed to cDNA using the SuperScript III reverse Transcriptase (Invitrogen) and random hexamers. DNA and cDNA extracts were quantified with a Qubit 1.0 (Thermo Fisher Scientific) and kept at −80°C.

### Sequencing and processing of picoeukaryotic community

Picoeukaryotic diversity was assessed by amplicon sequencing of the V4 region of the 18S rDNA gene (~380 bp) using the Illumina MiSeq platform and paired-end reads (2×250 bp). PCR amplifications with the eukaryotic universal primers TAReukFWD1 and TAReukREV3 (Stoeck et al 2010) and amplicon sequencing were carried out at the Research and Testing Laboratory (Lubbock, USA; http://www.researchandtesting.com). Illumina reads obtained from DNA and cDNA extracts (rDNA and rRNA sets, respectively) were processed together following an in-house pipeline (Logares 2017) at the Marine Bioinformatics Service (MARBITS) of the Institut de Ciències del Mar. Briefly, raw reads were corrected using BayesHammer (Nikolenko et al 2013) as indicated by Schirmer et al. (Schirmer et al 2015). Corrected paired-end reads were subsequently merged with PEAR (Zhang et al 2014) and sequences longer than 200 bp were quality-checked and dereplicated using USEARCH (Edgar 2010). OTU (Operational Taxonomic Unit) clustering at 99% similarity was done using UPARSE v8 (Edgar 2013). Chimera check and removal was performed both de novo and using the SILVA reference database (Quast et al 2013). Taxonomic assignment was done by BLAST searches against PR2 (Guillou et al 2013) and two in-house marine protist databases (available at https://github.com/ramalok) based on a collection of Sanger sequences from environmental surveys (Pernice et al 2013) and of 454 reads from the BioMarKs project (Massana et al 2015). Metazoan, Charophyta and nucleomorphs OTUs were removed. The final OTU table contained 79 rDNA samples and 90 rRNA samples (some samples were excluded due to suboptimal PCR or sequencing) accounting for 11,570,044 reads clustered into 45,115 OTUs. To enable comparisons between samples, the OTU table was randomly subsampled down to the minimum number of reads per sample (22,379 reads) using the *rrarefy* function in the *Vegan* package (Oksanen et al 2015). This turned into a final table including 38,300 OTUs and 3,782,051 reads.

### Statistical analyses

Statistical analyses were performed in R (R Core Team 2015) using the package *Vegan* (Oksanen et al 2015). Bray-Curtis dissimilarities were used as an estimator of beta diversity, which were then clustered using non-metric multidimensional scaling (NMDS). In NMDS, the differences between predefined groups were statistically tested with ANOSIM using 1,000 permutations. PERMANOVA analyses were performed to determine the proportion of the variation in community composition that was explained by the measured environmental variables. The Shannon index (H’) and richness (number of OTUs) were calculated as estimators of alpha diversity.

To assess the relative activity of taxonomic groups, rRNA:rDNA ratios were calculated for each OTU in each sample by dividing RNA between DNA reads. OTUs occurring in only one of the two datasets were not considered. Each individual ratio provided an indication of the relative activity of a given OTU in a given sample. Specifically, ratios >1 indicated metabolically “*hyperactive*” cells, ratios <1 indicated metabolically “*hypoactive*” cells, while ratios ~1 pointed to cells with “*average*” activity levels. Ratios from the same taxonomic group and water layer were analyzed together. Differences in rRNA:rDNA ratios by water layer were evaluated using a Wilcoxon test. For groups featuring predominantly heterotrophic species (therefore able to thrive in the dark), we analyzed the metabolic differences among the four layers, whereas for groups including mostly phototrophic species, we only focused in the surface and DCM layers (although they were sometimes detected in aphotic layers).

Some mesopelagic samples were taken in the oxygen minimum zone (OMZ) or in the deep scattering layer (DSL). For some analyses, DSL samples (accounting for 9 samples) were compared with the other mesopelagic samples (22 samples). DSL was identified by acoustic data (Irigoien et al 2014), and DSL samples were carefully selected from other mesopelagic samples according to their scattering profile. OMZ samples (8 samples) were considered where oxygen concentration was below 2 mg O_2_ L^-1^ (Vaqué-Sunyer and Duarte 2007), and were also compared with the rest of mesopelagic samples (23 samples).

## RESULTS

### Community structure along the water column

Alpha diversity estimates (richness and Shannon index) indicate that the diversity of picoeukaryotic assemblages decreased with depth in both rRNA and rDNA datasets (Fig. 1). The differences in richness between the epipelagic and the deep ocean layers were significant (Wilcoxon test p<0.05), with median values around 2,500 OTUs in the photic layer and 800 OTUs in the bathypelagic for both rRNA and rDNA datasets, while differences between Shannon indices were not significant. The same general pattern was observed when analyzing separately the Atlantic, Indian and Pacific oceans (Fig. S2), with higher diversity in the Pacific rRNA mesopelagic layer compared with that in other oceans.

**Fig. 1.**
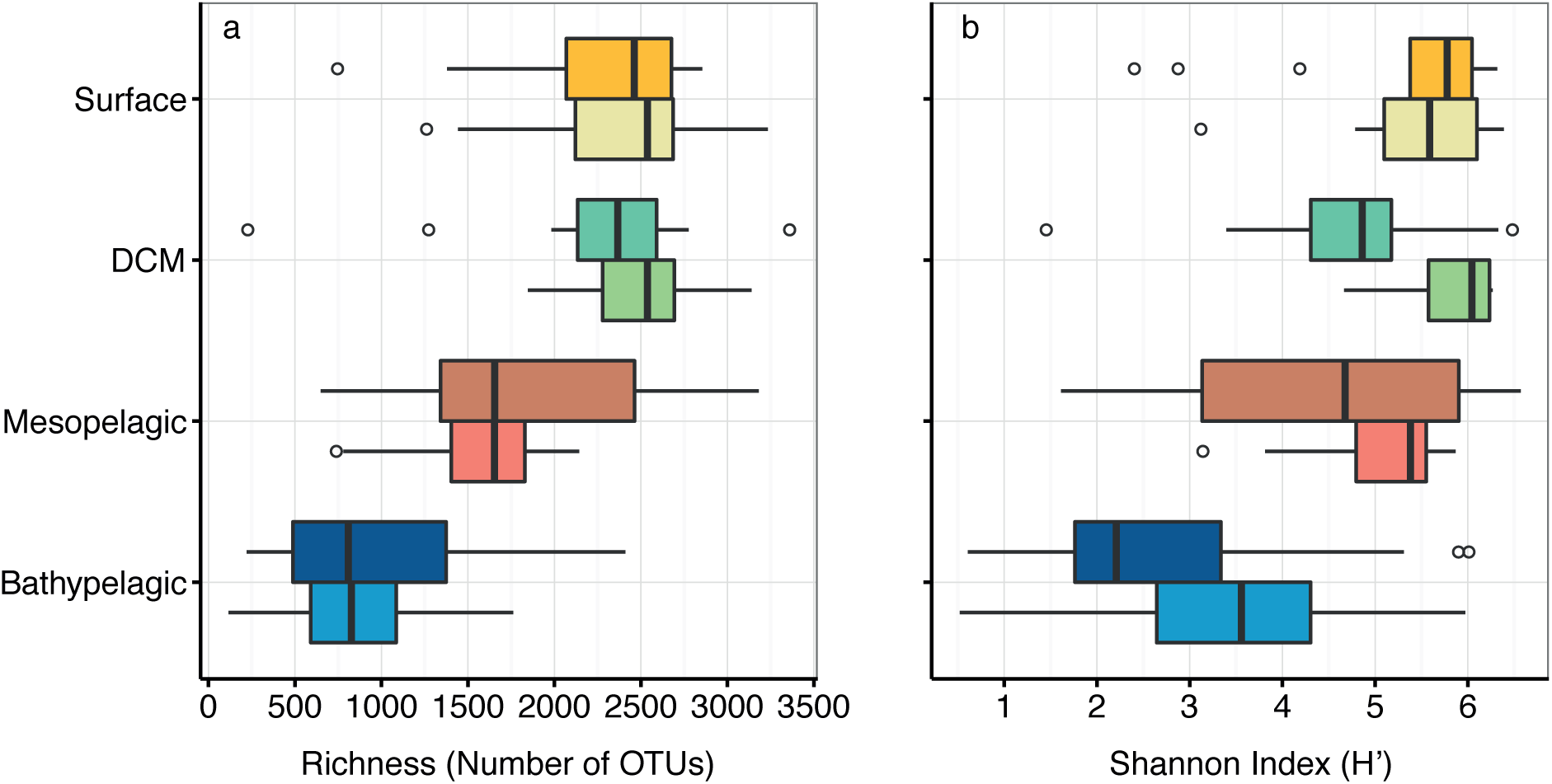
Alpha diversity of picoeukaryotic communities in the different water layers using the rRNA (upper boxplots) and rDNA (lower boxplots) datasets. (a) OTU richness and (b) Shannon Index (H’). Significant differences were found between photic and aphotic layers.

Community structure estimated from rDNA and rRNA formed two different clusters in the NMDS, featuring two groups, photic and aphotic communities, within each one (Fig. S3). This indicated that different community compositions were estimated by rDNA and rRNA extracts. Different NMDS for rDNA and rRNA datasets supported a significant differentiation between photic (surface and DCM) and aphotic (meso- and bathypelagic) communities (ANOSIM test: R^2^=0.52, p<0.001 for rRNA and R^2^=0.72, p<0.001 for rDNA, Fig 2). Within the photic layer, surface and DCM communities formed two groups (ANOSIM: R^2^=0.39 for rRNA and R^2^=0.60 for rDNA, p<0.001), while the mesopelagic and bathypelagic communities did not constitute different groups (ANOSIM test: R^2^=0.14 and R^2^=0.35, p<0.004, Fig. 2). Picoeukaryote communities from the Indian and Pacific oceans within each water layer formed separated clusters, while Atlantic communities were intermixed within the Indian and Pacific communities (Fig. 2).

**Fig. 2.**
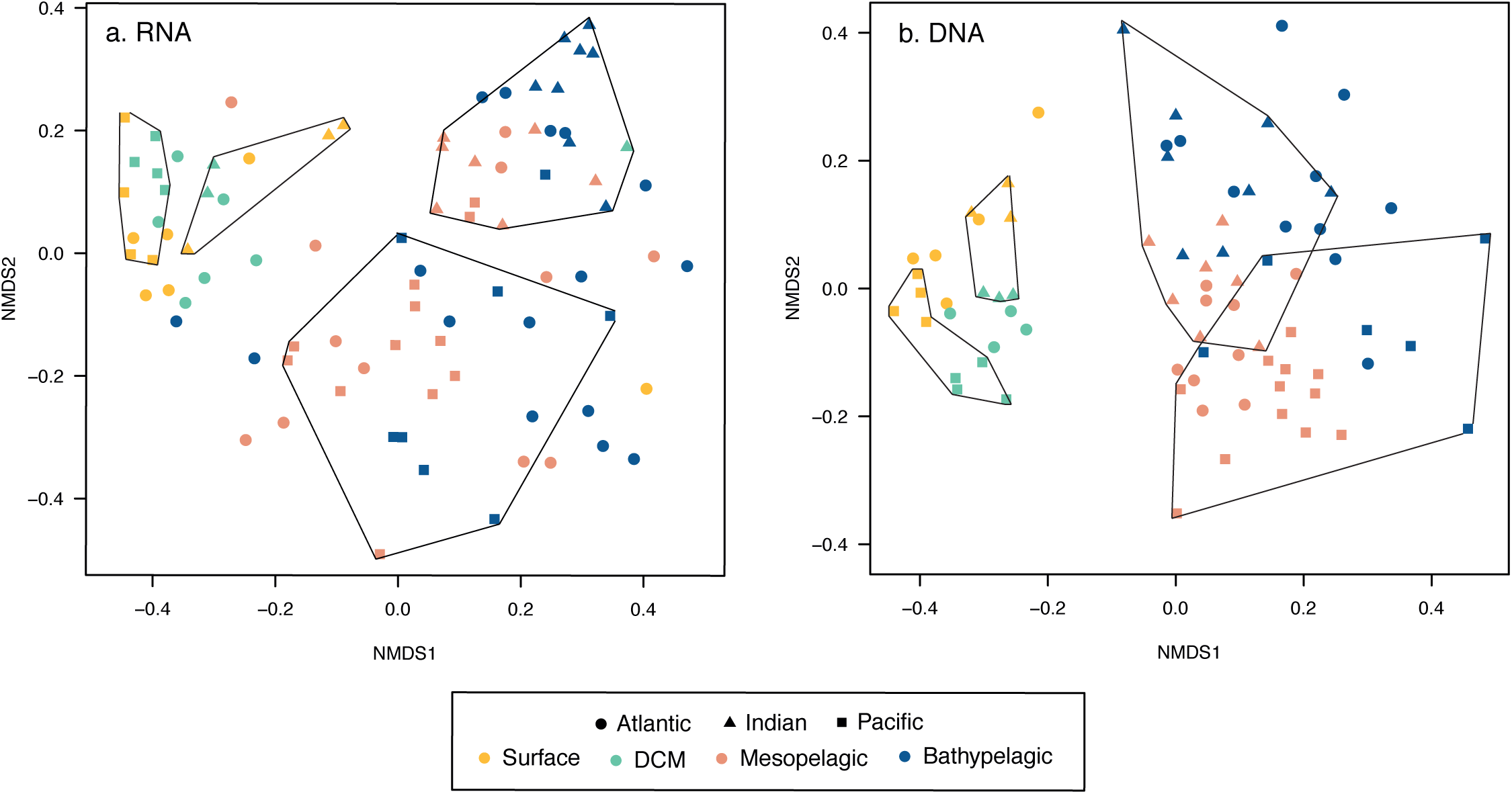
Comparison of picoeukaryotic community structure derived from the rRNA (a) and rDNA (b) datasets based on a non-metric multidimensional scaling analysis (NMDS). Each sample is colored according to the depth layer and has a different symbol shape according to the ocean where it comes from. Photic and aphotic samples from the Indian and the Pacific oceans were grouped in separate polygons.

Environmental parameters changed markedly along the water column (Fig. S4), likely exerting environmental selection. The variance in community structure that is explained by the measured environmental variables was analyzed with PERMANOVA. Light (taken as presence or absence) was the most important environmental factor structuring picoeukaryotic community, accounting for about 15% of community variance in both rRNA and rDNA datasets, while temperature, oxygen, the ocean basin (Atlantic, Indian, or Pacific), and depth explained each 3-7% of community variance along the water column. About 55-60% of the variance remained unexplained by the measured parameters (Table S1). Individual PERMANOVA tests for each water layer indicated that environmental factors explain 65% of the variance in community structure in the photic layer, whereas water mass explained about 25% of community structure variance in the dark ocean (Table S1).

### Horizontal community structure

We analysed the change in community composition within specific depth layers, which provides an indication of the connectivity between communities. We compared one sample of each water layer (i.e. 6 layers in rRNA and 5 in rDNA, as mesopelagic and bathypelagic samples were divided into two sublayers) among all stations (13 stations in rRNA and 10 in rDNA). Communities within the photic-zone, both from surface and DCM, were more similar among stations (Bray-Curtis distance 0.7, Figs. 3a, 3c) than those in the dark ocean, with those in the deepest bathypelagic layer sampled being the most dissimilar among stations (median Bray-Curtis values ~0.9). Interestingly, community composition in the bathypelagic was highly heterogeneous, as shown by the wide range of Bray-Curtis dissimilarities (ranging from 0.1 to 1.0). Most of the OTUs within each layer were found in a single station (Figs. 3b, 3d), with a decline in the number of OTUs with increasing prevalence (number of stations where a given OTU was present). This decline was not equal for all layers, being steepest in the bathypelagic layers. For instance, 3.3% of surface-OTUs were present in at least 80% of the samples, but this was 10 times lower (0.23% of OTUs) for the bathylepagic zone. This pattern of decreasing prevalence with depth is consistent with the higher Bray-Curtis dissimilarity of bathypelagic communities. The most prevalent OTUs were also the most abundant ones (data not shown).

**Fig. 3.**
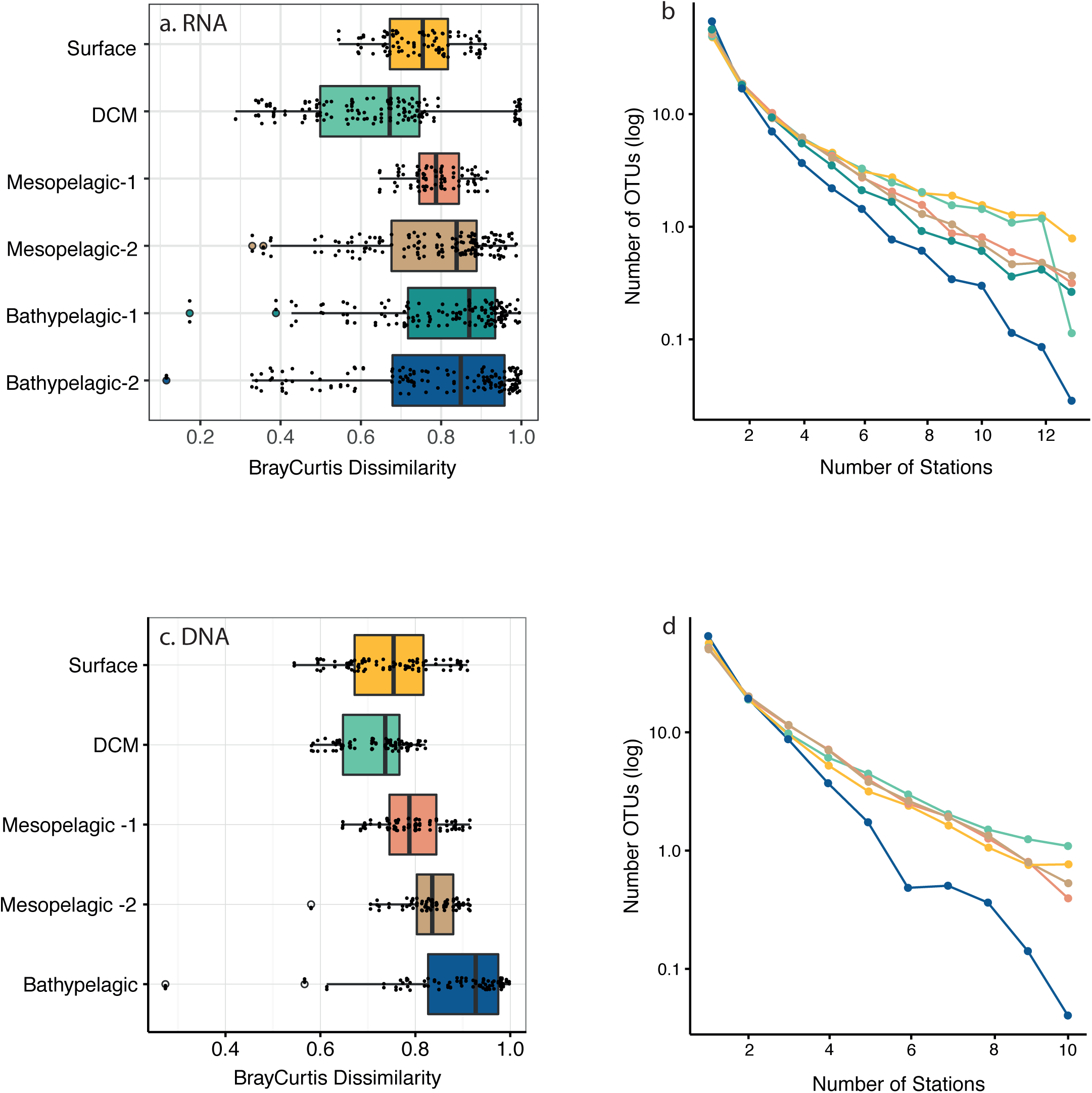
Community similarity and prevalence of OTUs across ocean depth layers, based on rRNA (upper panels) and rDNA (lower panels). (a, c) Distribution of Bray-Curtis dissimilarities values among samples from a given layer. (b, d) Number of OTUs within each water layer detected in a varying number of stations. The color lines correspond to the boxplot colors.

We also evaluated the potential vertical dispersal of OTUs in the water column. A total of 13% of OTUs (considering only those represented by >5 reads) in the rRNA dataset were restricted to the mesopelagic layer, compared to 4 to 6% of unique OTUs in the remaining depth layers (Table S2). A similar pattern was observed in the rDNA dataset, including a larger fraction of unique OTUs in the mesopelagic. Most shared OTUs were shared within the dark ocean (between mesopelagic and bathypelagic layers) or photic layers (between surface and DCM).

### Taxonomic change with depth

In general, there was a good correlation in the relative abundances of picoeukaryotic groups observed in the rDNA and rRNA datasets (Fig. S5). Striking exceptions were MALV-I, MALV-II and Polycystinea, highly overrepresented in the DNA survey (altogether 64.9% of rDNA reads but only 6.9% of rRNA reads) and Ciliophora, overrepresented in the rRNA survey (11.6% of rRNA reads versus 0.2% of rDNA reads). Acantharia, RAD-B, Mamiellophyceae and Basidiomycota were also overrepresented in the DNA dataset, while MOCH-5, Ancyromonadida and MAST-12 were overrepresented in the RNA dataset (Fig. S5).

We observed four patterns of taxonomic change with depth: (i) groups that increased their abundance with depth (Chrysophyceae, Bicosoecida and RAD-B), (ii) groups that decreased their abundance with depth (Dinoflagellata, Ciliophora, and all MAST and MOCH), (iii) groups that peaked in the mesopelagic (Labyrinthulomycetes, RAD-C, and MALV-IV) and (iv) groups that peaked at the DCM (Pelagophyceae, Mamiellophyceae, Telonema and Cryptomonadales) (Fig. 4). This resulted in group depth-dependent differences (based on rRNA), with Ciliophora and Dinoflagellata dominating in surface waters (42% of reads), Pelagophyceae and Dinoflagellata at the DCM (46% of reads) and Chrysophyceae and Bicosoecida in the dark ocean (40% of reads in the mesopelagic and 73% in the bathypelagic).

**Fig. 4.**
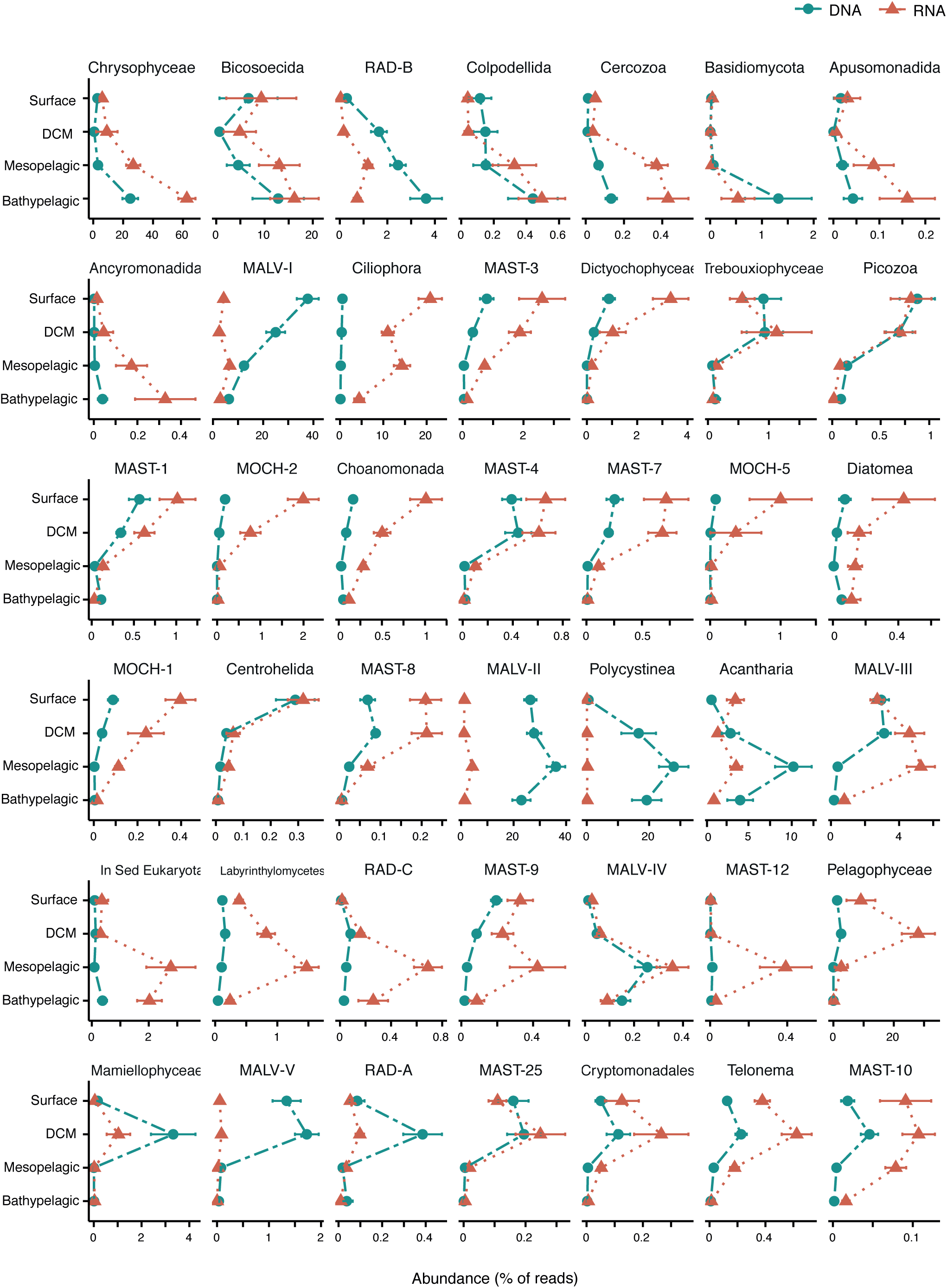
Averaged relative abundance of the main taxonomic groups in the four water layers derived from rRNA (red line) and rDNA (blue line) surveys. Groups are ordered first based on the layer of their maximal abundance, and then by their read abundance.

### Changes in the relative activity with depth

To determine the relative activity of OTUs, we calculated the ratio of rRNA vs. rDNA reads for all OTUs (31,866 ratios). The photic (surface, DCM) and mesopelagic layers had a median rRNA:rDNA ~1 for all OTUs, pointing to no deviations in relative activity (Fig. S6). In contrast the median rRNA:rDNA was < 1 in the bathypelagic zone (Fig. S6), pointing to a lower fraction of metabolically-active cells in deeper waters when compared to overlying waters. This lower relative activity in the bathypelagic was observed in the majority of heterotrophic groups (Fig. 5), which in general displayed the lowest activity in the bathypelagic (except MALV-I, Cercozoa, Labyrinthulomycetes, RAD-B and Telonema that showed the lowest activity at the photic zone). Many groups displayed higher relative activity in the mesopelagic (MALV-I, MALV-III, MALV-II, Cercozoa, Labyrinthulomycetes, among others, Fig. 5). Most phototrophic groups did not show significant differences in their relative activity between the surface and DCM layers, except for Prasinophyceae and MOCH-2 that displayed higher relative activity in the DCM (Fig. 5). No major differences were found in relative activity among ocean basins, although activities tended to be higher in the Pacific Ocean than in the Indian Ocean, specially for the aphotic layers (data not shown). Overall, relative activities were more variable with depth than among ocean basins.

**Fig. 5.**
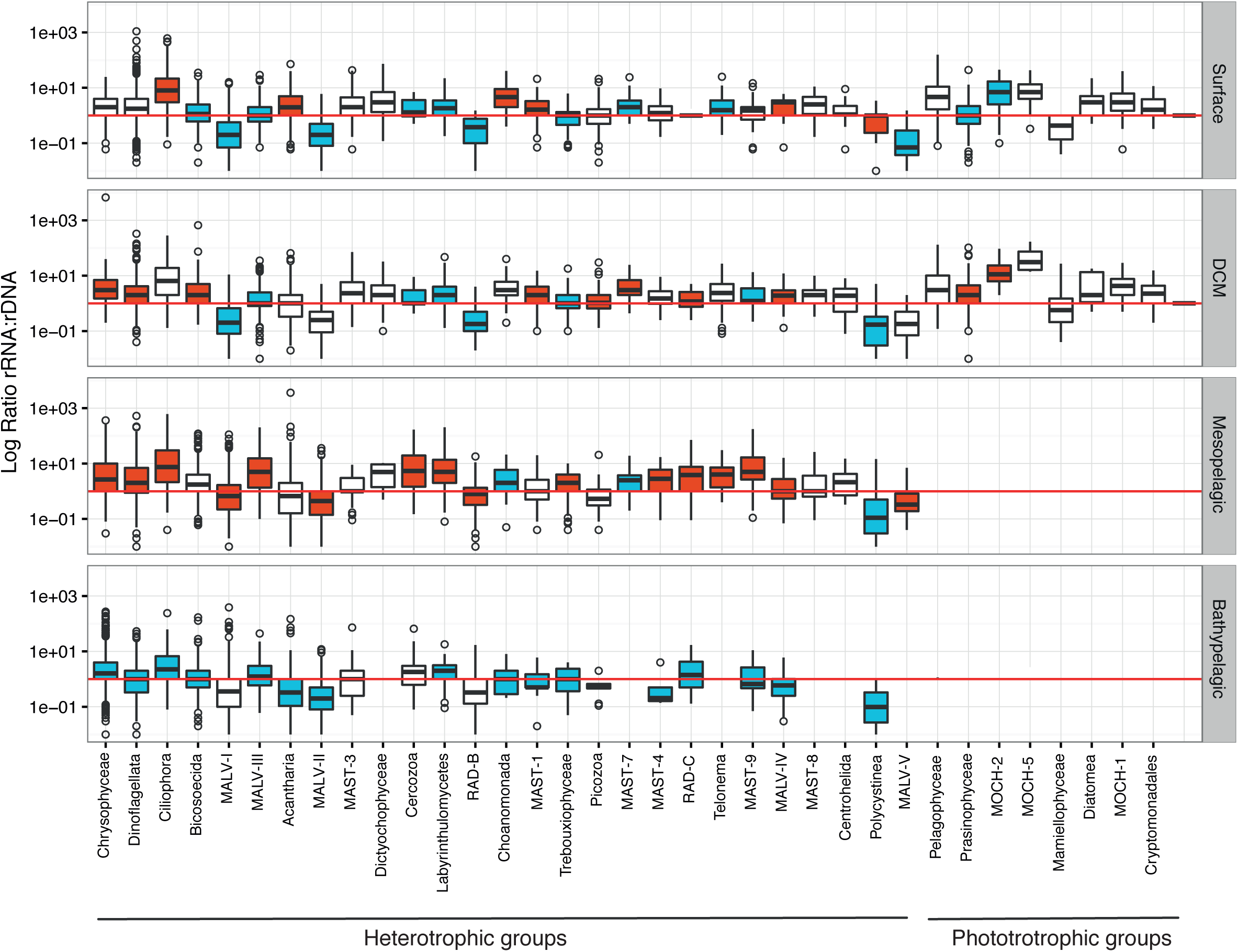
Distribution of the activity ratios (rRNA:rDNA) for all OTUs within major taxonomic groups in the four investigated depth layers. The red line indicates a ratio of 1. For each group, the layers with significantly higher and lower values between them, according to Wilcoxon tests, are colored in red and blue, respectively. Groups with no boxplot color (e.g. MAST-3), indicate no significant differences among layers. Note that heterotrophic taxa are listed first, and then the phototrophic ones.

### Patterns in OMZ and DSL

We sampled the OMZ and DSL in specific stations (eight samples of 3 Pacific Ocean stations were sampled in the OMZ, and DSL was sampled in 9 of the 13 stations). Whereas no clear differentiation was observed between DSL and the remaining mesopelagic communities (Fig. S7a), OMZ communities were more similar among themselves and differed significantly from the remaining mesopelagic samples (Fig. S7b). The DSL communities did not display a different richness than the rest of the stations, while the OMZ had a higher richness than the remaining mesopelagic samples (Fig. S8).

We explored which taxonomic groups preferred the OMZ as compared with the fully oxygenated mesopelagic waters (Fig. 6a). Ciliophora, Dinoflagellata, MALV-III, MALV-II and Acantharia were enriched in the OMZ, accounting for 45% of the total mesopelagic reads. Groups enriched in the OMZ tended to have higher activities (rRNA:rDNA ratio) in that region, except for Chrysophyceae and Bicosoecida, which were more active but less abundant in the OMZ (Fig. 6a). Two OTUs with contrasting behavior explained this difference in the Crysophyceae, one being very abundant in the OMZ and the other in the mesopelagic, whereas no clear pattern was found for Bicosoecida. The group formed by unclassified OTUs (IncertaeSedis Eukaryota), which could potentially include new species within high-rank taxa, showed a higher abundance and relative activity in the OMZ, pointing to potential taxonomic novelty. A similar analysis of DSL samples yielded inconclusive results (Fig. 6b). Some taxonomic groups were enriched in the DSL (MALV-I, Pelagophyeae, MAST-9), but the relative abundance ratio was, in general, smaller for most groups, indicating no clear DSL preference. Surprisingly, most groups showed lower activities at the DSL, suggesting that the DSL is not a specific and selective habitat for picoeukaryotes, in contrast with the OMZ that influences picoeukaryotic community structure and induces changes in their relative activity.

**Fig. 6.**
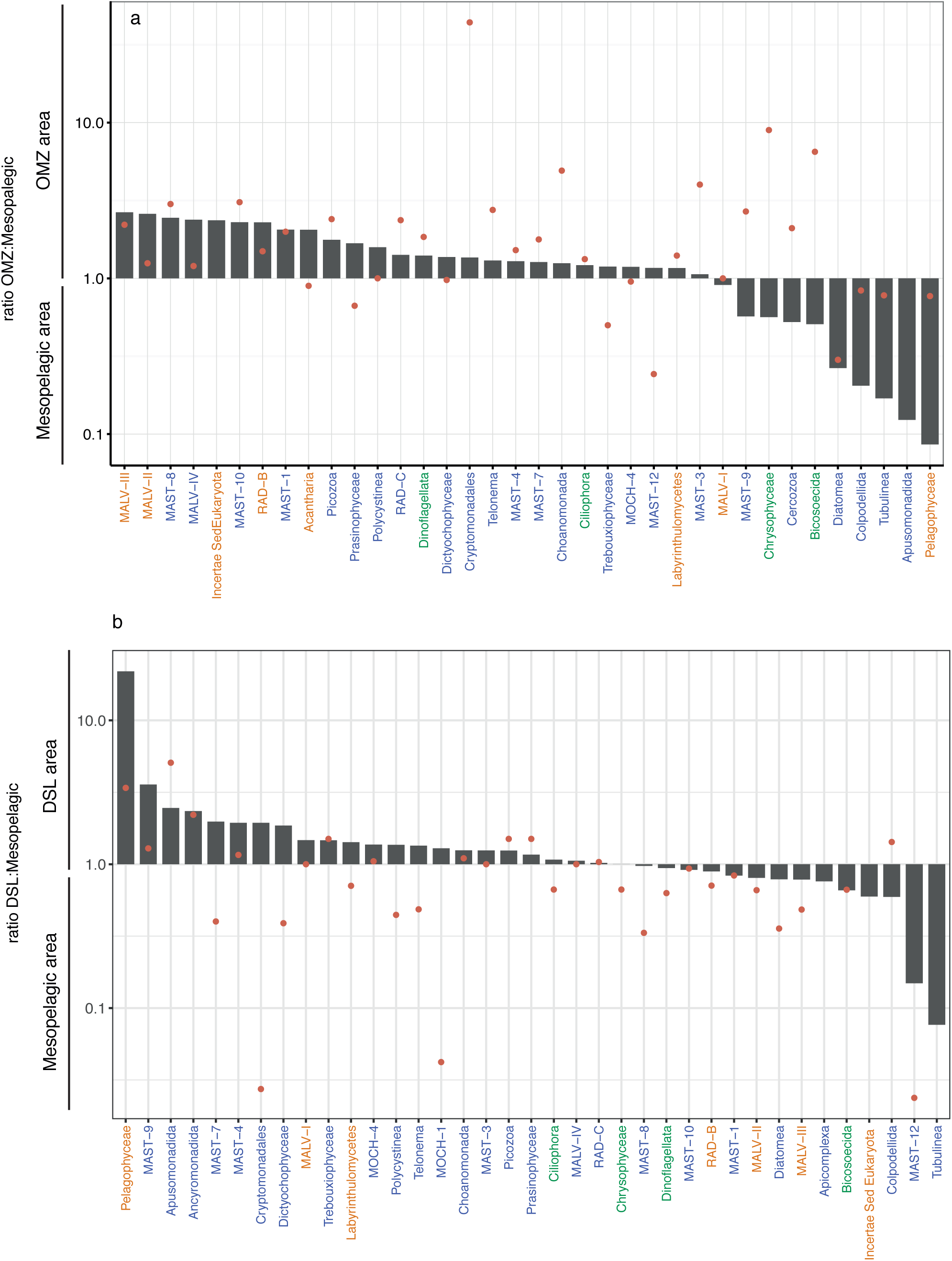
Comparison of relative abundance of taxonomic groups and their relative activity in the communities sampled within the Oxygen Minimum Zone (a) and DSL (b) with respect to the rest of mesopelagic samples. Bars represent the ratio between the abundance of specific groups among the two regions, while dots represent the ratio of the relative activity (rRNA:rDNA). Groups are ordered by their higher abundance ratio in the OMZ region (a) and DSL region (b) (only groups with an abundance >0.05% in the total mesopelagic realm are shown). Note that bars or dots in the OMZ zone of the plot mean higher abundance or activity in that zone. Group colors represent the relative abundance of the group in the total mesopelagic: abundance >10% in green, 1-10% in yellow, and 0.5-1% in blue.

## DISCUSSION

Our work provides the first global assessment of the change in community structure and relative activity of picoeukaryotes from surface down to 4,000 m. We found clear patterns of diversity and relative activity change along the water column, with clear differences in picoeukaryotic assemblages between the photic and aphotic regions, consistent with results from previous regional surveys (Brown et al 2009, Countway et al 2007). Furthermore, different picoeukaryotic communities were observed in surface and DCM samples within the photic region, and between mesopelagic and bathypelagic samples within the aphotic region. Light, temperature and inorganic nutrients were found to drive community structure in surface and DCM layers, but not in the dark ocean. The mesopelagic layer contained the highest number of unique picoeukaryotic OTUs, in agreement with previous regional observations (Brown et al 2009). This is partially attributable to the existence of waters with specific conditions, such as the oxygen minimum zone. Oxygen concentration has a strong influence on microbial distributions, with only a subset of taxa able to adapt to anoxic or microoxic conditions (Orsi et al 2012). Despite the higher number of unique OTUs in the mesopelagic, this layer was not the most diverse, as richness was higher in surface waters and decreased with depth, which is coincident with previous studies (Brown et al 2009, Countway et al 2007). This is, in part, expected as obligate phototroph are restricted to photic layers.

The abundance of the different picoeukaryotic taxonomic groups changed with depth, with Chrysophyceae, Bicosoecida, Radiolaria and Colpodellida increasing in relative abundance with depth, as reported in earlier regional studies (Brown et al 2009, Countway et al 2007, Edgcomb et al 2002, Hu et al 2016, Not et al 2007). RAD-C was an important component of twilight and dark deep-ocean communities, showing a peak in relative abundance in the mesopelagic layer. Radiolaria are typically very large and sometimes fragile protists, so they can break during size fractionation, but can also produce swarmers of picoplankton size, so their detection in picoplanktonic studies is still controversial.

The abundance of photosynthetic groups (e.g. Pelagophyceae, and green algae) declined, as expected, with depth, as well as several heterotrophic lineages such as MAST clades or Picozoa. The occasional detection of metabolically active phototrophic groups in the deep-ocean, which have already been detected sometimes at significant abundances (Agustí et al 2015, Xu et al 2017), could be due to their attachment to rapidly sinking particles (Agustí et al 2015), although some of the detected taxa may be mixotrophs, as this lifestyle is thought to be more common in the ocean than currently acknowledged (Unrein et al 2014).

Even though a moderate correlation of rDNA and rRNA relative abundances was found for most taxonomic groups, our results indicated that some groups were overrepresented in the rRNA dataset, whereas others like MALV-I, MALV-II, Polycystinea, Acantharia were overrepresented in the rDNA dataset. A general explanation for this pattern would be that the latter groups have a relatively higher rDNA copy number than the rest of the community. MALV-I and MALV-II are usually dominant groups in rDNA surveys (Bachy et al 2011, de Vargas et al 2015, López-García et al 2001, Massana et al 2015, Pernice et al 2016) whereas they may be up to ten times less abundant in rRNA surveys (Massana et al 2015). It is accepted that they likely have many rDNA operon copies (Siano et al 2010) indicating that, in most cases, rDNA-based assessments may overestimate their abundance. In a previous study we demonstrated that metabarcoding based on DNA and RNA extracts provide reasonable views of relative abundance, with different results depending on the taxonomic group (Giner et al 2016), indicating that the patterns observed here are robust.

Epipelagic communities were more similar among themselves across the ocean than communities in the dark ocean, suggesting a higher dispersion in surface and DCM communities, likely due to currents in the upper ocean (Villarino et al. 2018). Bathypelagic assemblages, which were the most different across the ocean (Fig. 3), seemed to be structured by water masses, which accounted for 32% of the variability in their community structure. Thus, two distinct water-masses, even geographically close, may contain different communities, and vice versa (Pernice et al 2016). More OTUs where shared between the two deep-ocean layers than between the two epipelagic. We hypothesize that dispersion could be limited by environmental conditions and the adaptability of the microorganism. So, when a given taxa is adapted to an environmental constrain like the lack of light and high pressure, then it can be found in both deep layers.

The comparison among rDNA and rRNA datasets provide insights on group relative activity, which represents, in addition to the global scale of our study, an innovative aspect of the research presented here, as most picoeukaryotic surveys so far have been based in rDNA gene from DNA extracts, while few studies have also included rRNA extracts (Hu et al 2016, Logares et al 2014, Massana et al 2015, Not et al 2009, Xu et al 2017). However, comparisons of rRNA:rDNA ratios among different taxa may be confounded by the potentially large variation of rDNA copy number among taxonomic groups (Prokopowich et al 2003), and cells of different sizes (Zhu et al 2005). Still, within a particular taxonomic group, the rRNA copy number will vary depending on the relative activity and, in contrast to rDNA, rRNA is assumed to be absent from the extracellular pool (Blazewicz et al 2013). In our analyses, many heterotrophic groups showed their highest relative activity in the mesopelagic layer, where oxyclines may promote hotspots of metabolic activity (Edgcomb 2016), and large carbon transfer by migratory fish and invertebrates may promote relatively high prey abundance to be grazed by metabolically active communities of Ciliates, Dinoflagellates and Cercozoans, which were some of the most active groups in the mesopelagic layer (Fig. 6). Furthermore, it has been observed that the clearance rates of heterotrophic nanoflagellates were higher in mesopelagic than in epipelagic samples (Cho et al 2000). Relative activities of Cercozoa, MALV-I, MALV-II were also high in the mesopelagic, in agreement with observations by Xu et al (2017). Also, Hu et al. (2016) found higher metabolic activity of Ciliophora at those depths. On the other hand, as expected, the majority of taxa were less active in the bathypelagic. Our analysis found depth-related patterns in the relative activity for most of the groups and shed new light on relatively unknown groups, such as the MALV-III, and highlight the potential important role of protists in the deep ocean.

Low oxygen conditions, often found within the mesopelagic layer, can affect the community structure of protists (Jing et al 2015, Orsi et al 2012, Parris et al 2014, Wylezich et al 2018). Yet, protist distributions in OMZ are largely unknown. Our results indicated that the alpha diversity found in OMZ in the Pacific Ocean is similar to that of epipelagic waters, as observed by Jing et al (2015), indicating a relatively high diversity in hypoxic waters. The most abundant groups in the OMZ retrieved with the rRNA were Chrysophyceae, Ciliates and Dinoflagellates. The presence of Chrysopyceae in the OMZ has been reported before (Jing et al 2015), together with that of anaerobic Ciliates (e.g. mesodiniids) and Dinoflagellates (Orsi et al 2012, Parris et al 2014, Wylezich et al 2018). The OMZ contains a diverse community of picoeukaryotes, with 25 different taxonomic groups present, most of them showing higher relative abundances (Fig. 6). Furthermore, the fact that most of the taxonomic groups were metabolically active could be explained by the high bacterivory activity of mixotrophic and heterotrophic groups, as bacterial abundance is typically high below the oxycline (Jing et al 2015, Fenchel and Finlay, 2008), where ciliates are recognized as important grazers (Parris et al 2014). Besides, species richness was higher in the OMZ than in the mesopelagic zone, a pattern previously observed by Parris et al (2014) but that differs from other reports (Orsi et al 2012). Yet, contrary to what we expected, we did not find conspicuous picoeukaryote assemblages in the DSL as compared with the rest of mesopelagic samples, and intriguing observation that deserves further explorations.

In conclusion, this study provides the first insights on the changes in diversity and relative activity of picoeukaryotes along the whole water column at the global scale. Picoeukaryotic community structure was strongly differentiated in the water column, with two main communities corresponding to the epipelagic and the dark ocean. Our analysis identified the mesopelagic layer as a diversity hotspot for picoeukaryotes, indicating also differentiated communities within oxygen minimum zones.

## ACKNOWLEDGEMENTS

This project was supported by the Spanish Ministry of Economy and Competitiveness through project Consolider-Ingenio Malaspina-2010 (CSD2008–00077). Further research was funded by Spanish projects FLAME (CGL2010-16304, MICINN), ALLFLAGS (CTM2016-75083-R, MINECO), INTERACTOMICS (CTM2015-69936-P, MINECO/FEDER) and REMEI (CTM2015-70340-R). CRG was supported by a Spanish FPI grant. We thank all the scientists that contributed samples and data for this study in the different legs of the Malaspina Expedition, as well as the crew of the R/V BIO Hespérides. Bioinformatic analyses have been performed at the Marbits platform (ICM-CSIC; https://marbits.icm.csic.es).

## CONFLICT OF INTEREST

The authors declare no conflict of interest.

