## Supplementary information for "Marked changes in diversity and relative activity of picoeukaryotes with depth in the global ocean"

#### SUPPLEMENTARY FIGURES

**Figure S1.** World map showing the location of the Malaspina stations sampled for this study. Orange dots represent sations containing Oxygen Minimum Zone (OMZ) samples.

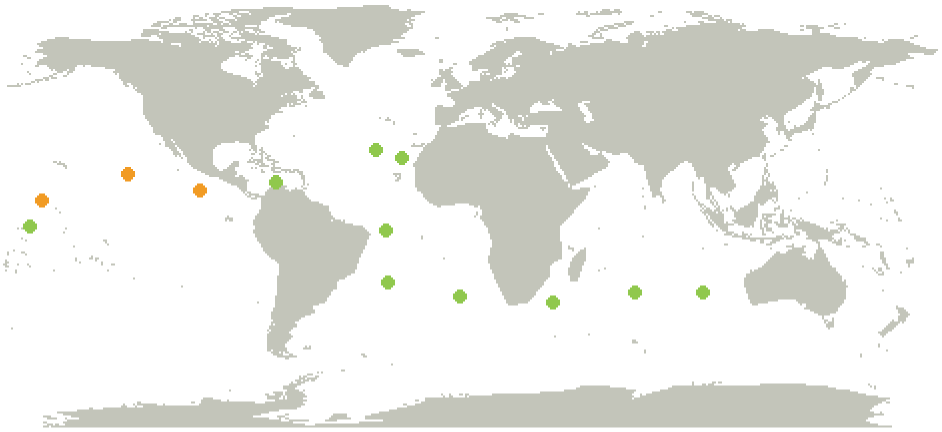

**Figure S2.** Richness of picoeukaryotic communities in samples from different water layers of each ocean basin using the rRNA (upper boxplots) and rDNA (lower boxplots) datasets.

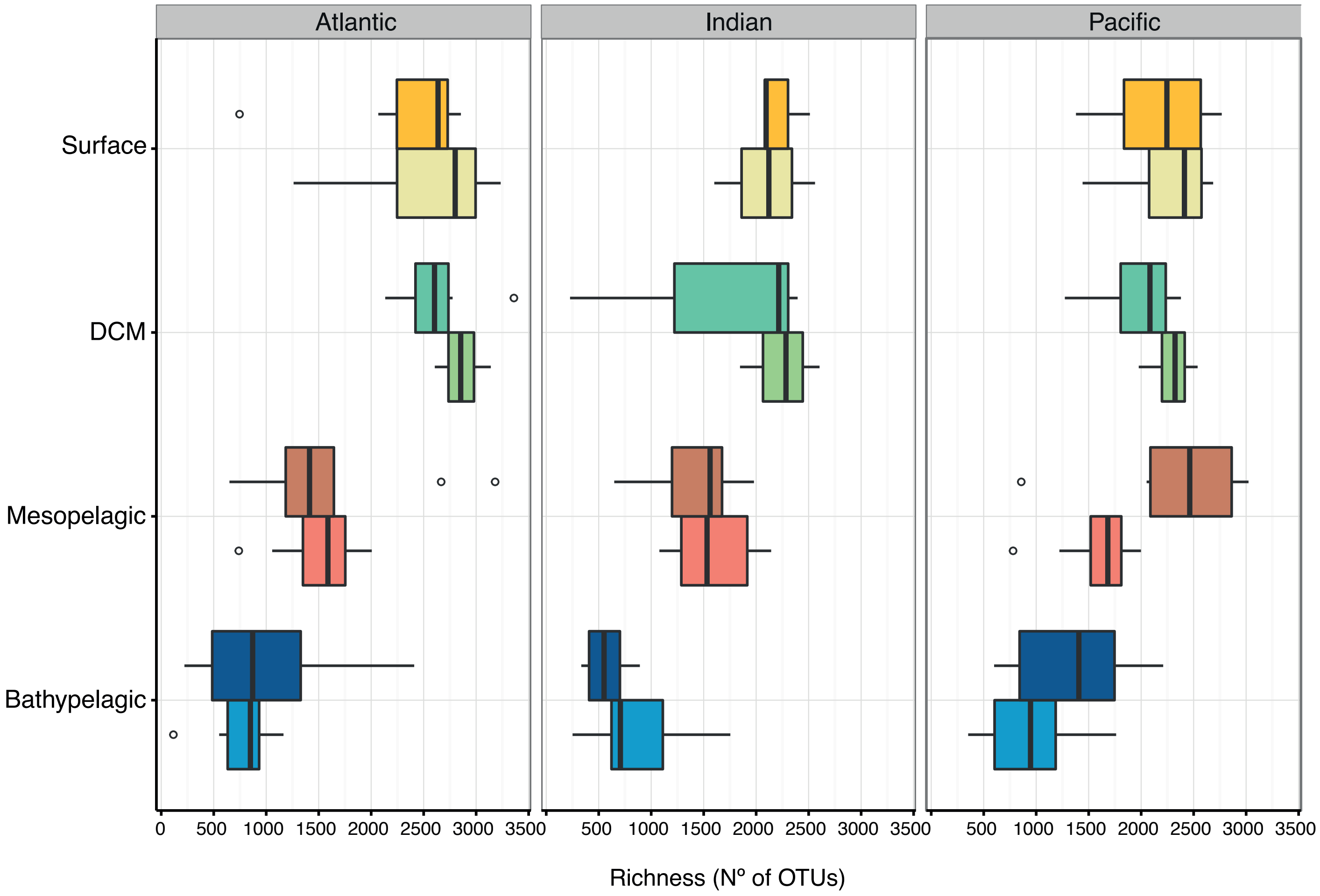

**Figure S3.** Clustering of all picoeukaryotic samples in a non-metric multidimensional scaling (NMDS) plot. Each sample is colored according to the depth layer and has different symbol shape according to the type of dataset (rDNA or rRNA).

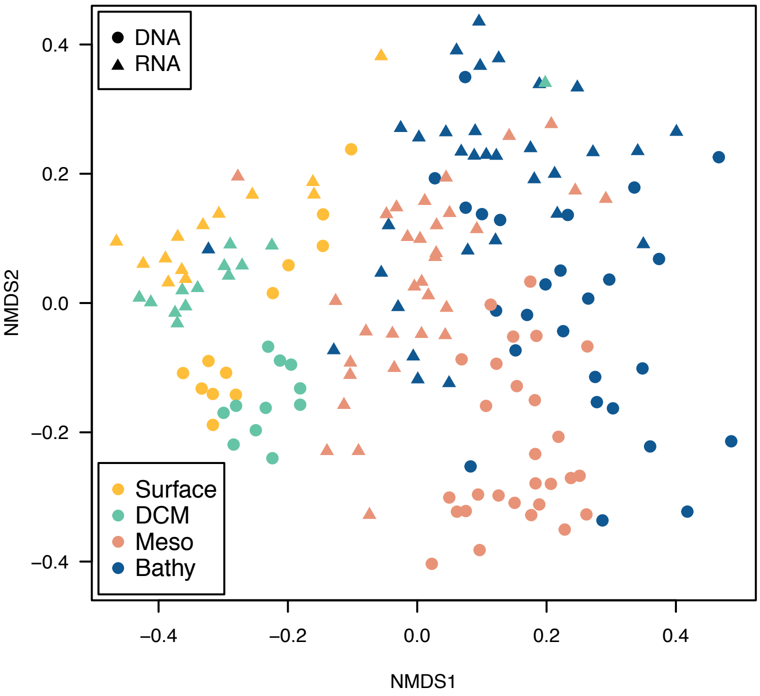

**Figure S4.** Main environmental variables and inorganic nutrient concentrations in the four layers of the water column (actual values as black dots and averaged values as brown lines).

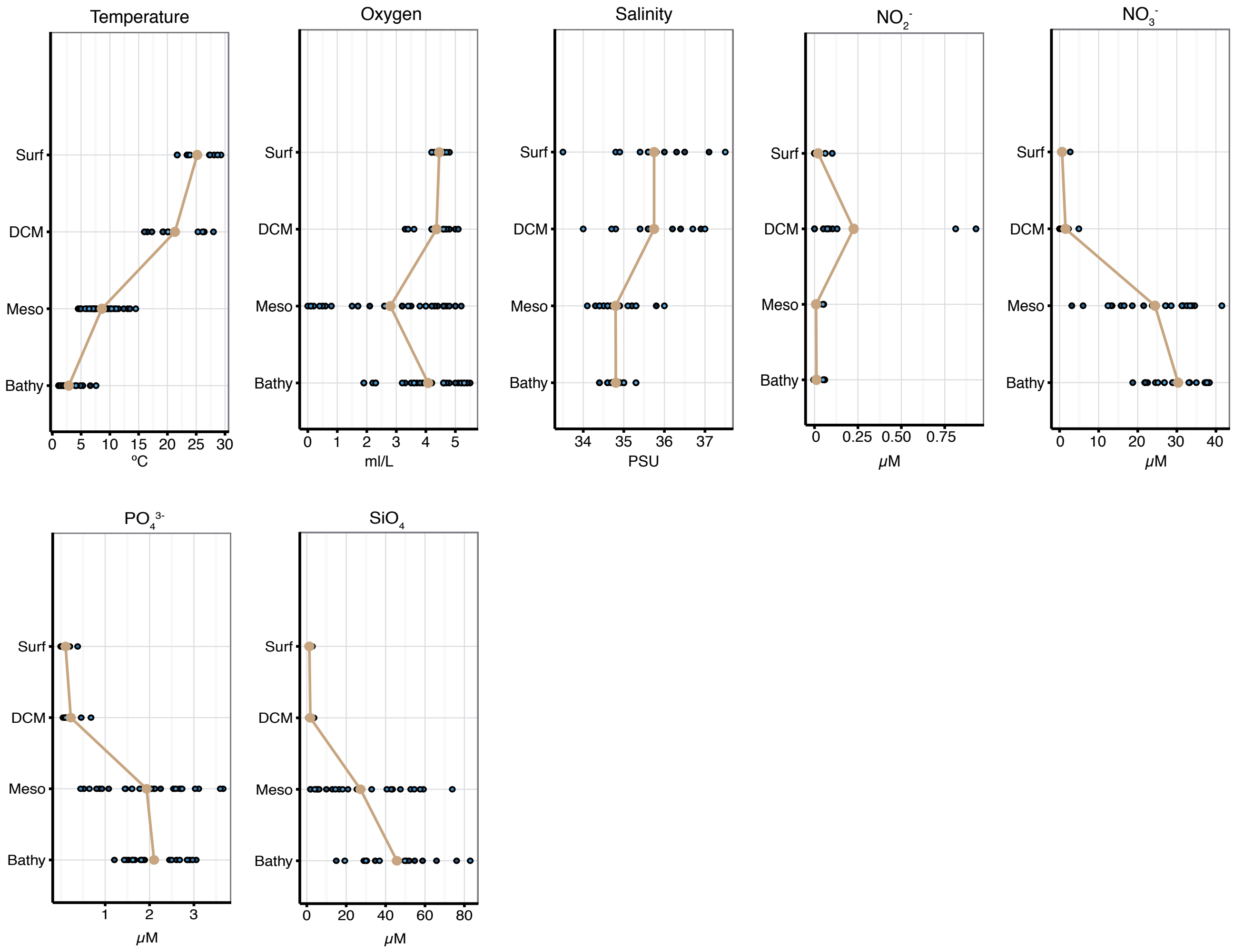

**Figure S5.** Representation of all phylogenetic groups based in the total abundance in the rRNA and rDNA surveys.

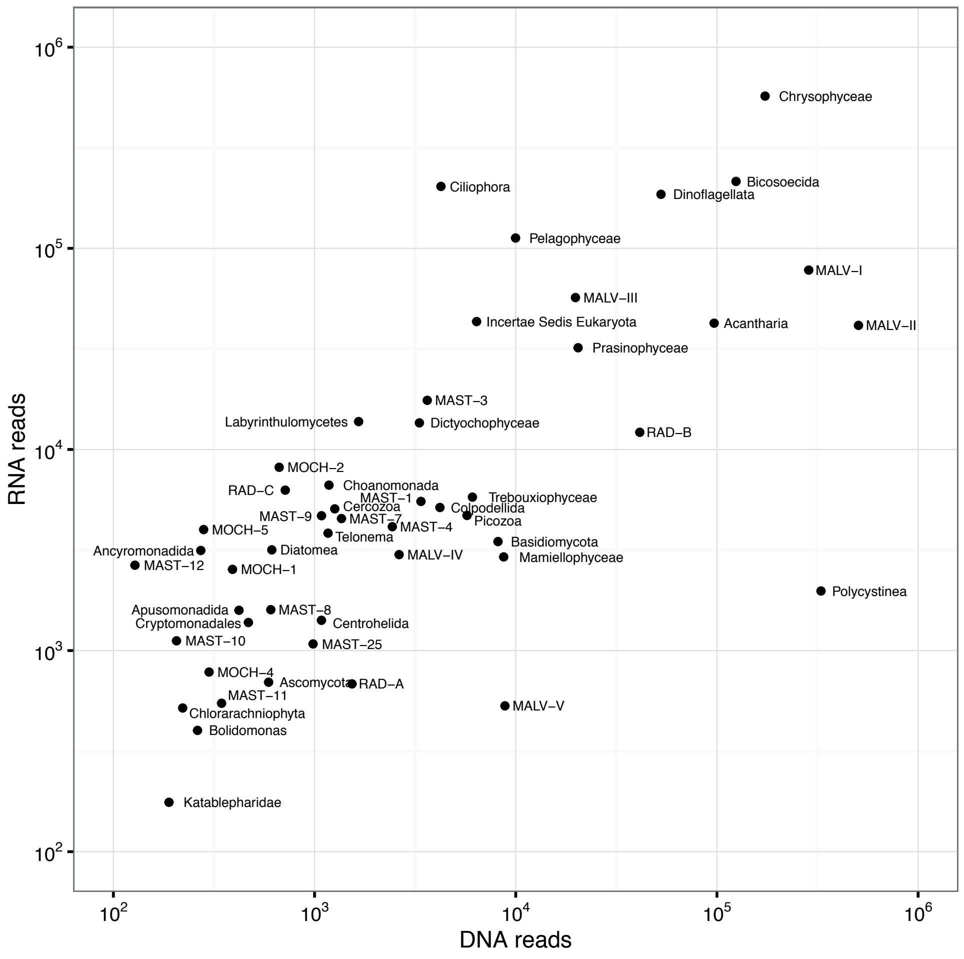

**Figure S6.** Distribution of the rRNA:rDNA ratios for all OTUs within a given depth layer. The red line indicates a ratio of 1.

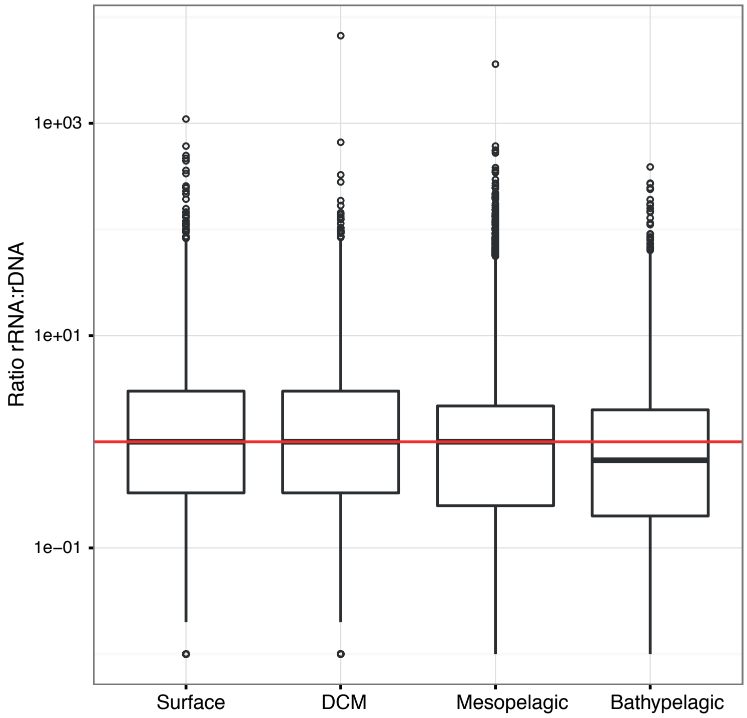

**Figure S7.** Clustering of all picoeukaryotic samples on a Non-metric multidimensional analysis (NMDS) based on Bray-Curtis dissimilarities differentiating between DSL (a, c) and OMZ (b, d) samples. Each sample is colored according the specific depth layer.

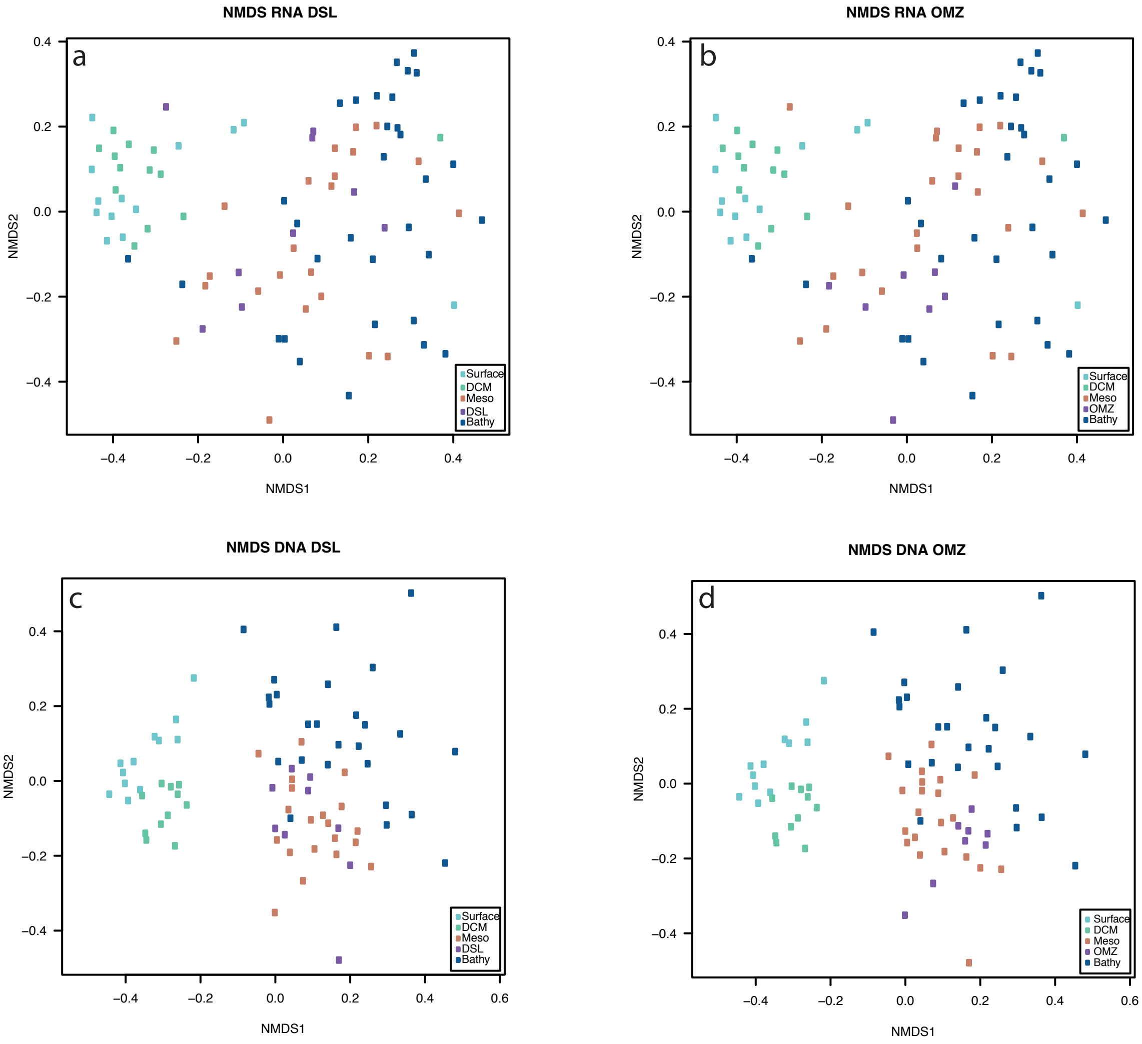

**Figure S8.** Richness in the different water layers defined as DSL (a) and OMZ (b). rRNA (upper boxplots) and rDNA (lower boxplots).

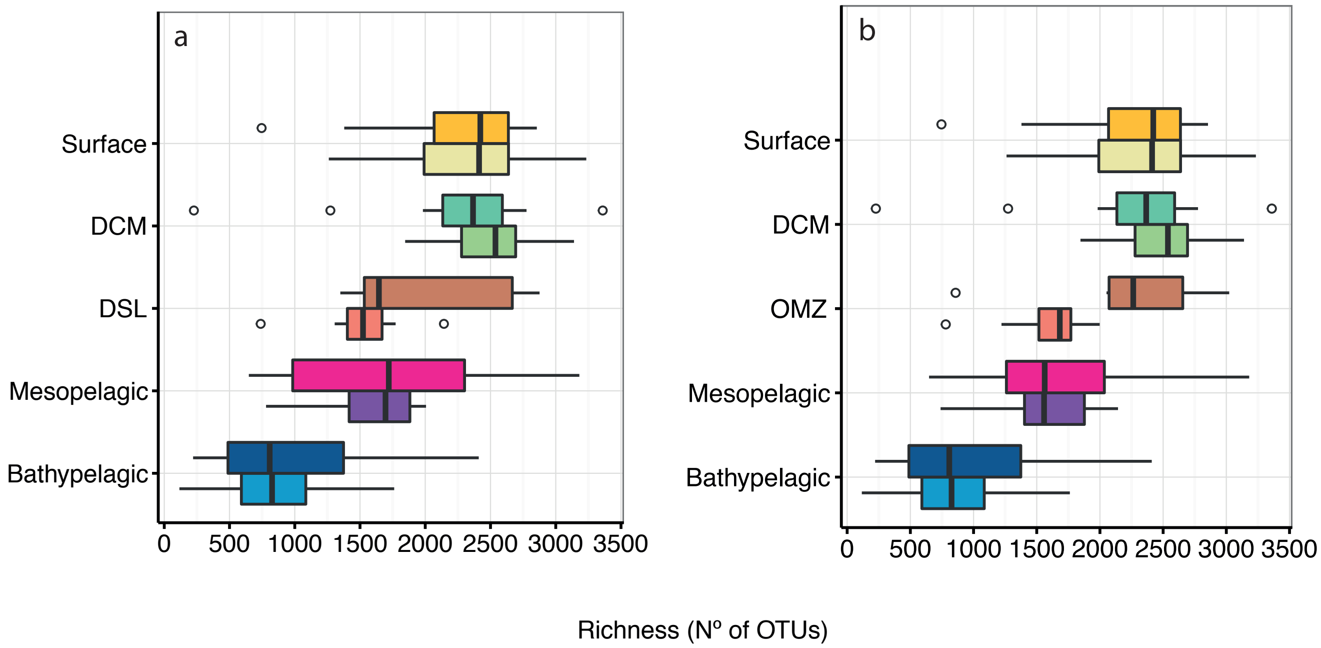

### SUPPLEMENTARY TABLES

**Table S1**. Results of the PERMANOVA analysis for the rDNA and rRNA datasets. For each dataset whole water column, and only epipelagic and deep-ocean depths had been analyzed.

| | DNA | | | | | |
|---|---|---|---|---|---|---|
| | **Whole Water column** | | **Surface + DCM (Epipelagic)** | | **Meso + bathypelagic (Deep – Ocean)** | |
| | **R2** | **p-value** | **R2** | **p-value** | **R2** | **p-value** |
| **Light (presence/absence)** | 0.14 | 0.001 | - | - | - | - |
| **Temperature** | 0.04 | 0.001 | 0.11 | 0.002 | 0.08 | 0.001 |
| **Salinity** | 0.01 | 0.139 | 0.06 | 0.082 | 0.03 | 0.021 |
| **Water Mass** | - | - | - | - | **0.24** | 0.045 |
| **$O_2$** | 0.04 | 0.001 | 0.07 | 0.014 | 0.06 | 0.001 |
| **Ocean** | 0.05 | 0.001 | 0.12 | 0.039 | 0.08 | 0.001 |
| **Depth** | 0.07 | 0.001 | 0.09 | 0.001 | 0.01 | 0.783 |
| **$NO_3$** | 0.01 | 0.511 | 0.04 | 0.374 | 0.02 | 0.275 |
| **$PO_4$** | 0.02 | 0.042 | 0.04 | 0.740 | 0.02 | 0.105 |
| **$SiO_4$** | 0.01 | 0.445 | 0.04 | 0.704 | 0.02 | 0.386 |
| **DAPI** | 0.01 | 0.122 | 0.05 | 0.268 | 0.01 | 0.864 |
| **Conductivity** | 0.02 | 0.017 | 0.04 | 0.495 | 0.03 | 0.017 |

| | RNA | | | | | |
|---|---|---|---|---|---|---|
| | **Whole Water column** | | **Surface + DCM (Epipelagic)** | | **Meso + bathypelagic (Deep – Ocean)** | |
| | **R2** | **p-value** | **R2** | **p-value** | **R2** | **p-value** |
| **Light (presence/absence)** | 0.15 | 0.001 | - | - | - | - |
| **Temperature** | 0.03 | 0.001 | 0.08 | 0.045 | 0.05 | 0.001 |
| **Salinity** | 0.02 | 0.017 | 0.09 | 0.032 | 0.04 | 0.023 |
| **Water Mass** | - | - | - | - | **0.25** | 0.040 |
| **$O_2$** | 0.06 | 0.001 | 0.04 | 0.296 | 0.02 | 0.173 |
| **Ocean** | 0.06 | 0.001 | 0.13 | 0.052 | 0.07 | 0.006 |
| **Depth** | 0.07 | 0.001 | 0.11 | 0.004 | | |
| **$NO_3$** | 0.01 | 0.227 | 0.03 | 0.777 | 0.01 | 0.989 |
| **$PO_4$** | 0.02 | 0.077 | 0.03 | 0.563 | 0.02 | 0.260 |
| **$SiO_4$** | 0.01 | 0.819 | 0.02 | 0.872 | 0.01 | 0.506 |
| **DAPI** | 0.01 | 0.268 | 0.08 | 0.038 | 0.02 | 0.317 |
| **Conductivity** | 0.01 | 0.343 | 0.04 | 0.438 | 0.02 | 0.320 |

**Table S1**. Number of total OTUs in each water layer and of the unique OTUs within them in the DNA and RNA dataset.

**DNA**

| Depth | Total OTUs | Nº of reads | OTUs unique | Nº reads unique | %reads unique |
|---|---|---|---|---|---|
| Surface | 5968 | 259,490 | 765 | 11,221 | 4.3 |
| DCM | 6741 | 239,147 | 849 | 16,989 | 7.1 |
| Mesopelagic | 8106 | 643,049 | 1652 | 47,084 | 7.3 |
| Bathypelagic | 6117 | 578,182 | 584 | 14,549 | 2.5 |

**RNA**

| Depth | Total OTUs | Nº of reads | OTUs unique | Nº reads unique | %reads unique |
|---|---|---|---|---|---|
| Surface | 6171 | 285,018 | 386 | 5,415 | 1.9 |
| DCM | 6891 | 285,525 | 284 | 3,339 | 1.2 |
| Mesopelagic | 9622 | 685,314 | 1221 | 22,967 | 3.35 |
| Bathypelagic | 8046 | 711,808 | 314 | 4,510 | 0.6 |
